# Structural and Functional Analysis of the D614G SARS-CoV-2 Spike Protein Variant

**DOI:** 10.1101/2020.07.04.187757

**Authors:** Leonid Yurkovetskiy, Xue Wang, Kristen E. Pascal, Christopher Tomkins-Tinch, Thomas Nyalile, Yetao Wang, Alina Baum, William E. Diehl, Ann Dauphin, Claudia Carbone, Kristen Veinotte, Shawn B. Egri, Stephen F. Schaffner, Jacob E. Lemieux, James Munro, Ashique Rafique, Abhi Barve, Pardis C. Sabeti, Christos A. Kyratsous, Natalya Dudkina, Kuang Shen, Jeremy Luban

**Author notes:** These authors contributed equally to this work (Leonid Yurkovetskiy, Xue Wang). Correspondence (P.C.S.), (C.A.K.), (N.D.), (K.S.), (J.L.).

## Abstract

The SARS-CoV-2 spike (S) protein variant D614G supplanted the ancestral virus worldwide in a matter of months. Here we show that D614G was more infectious than the ancestral form on human lung cells, colon cells, and cells rendered permissive by ectopic expression of various mammalian ACE2 orthologs. Nonetheless, D614G affinity for ACE2 was reduced due to a faster dissociation rate. Assessment of the S protein trimer by cryo-electron microscopy showed that D614G disrupts a critical interprotomer contact and that this dramatically shifts the S protein trimer conformation toward an ACE2-binding and fusion-competent state. Consistent with the more open conformation, neutralization potency of antibodies targeting the S protein receptor-binding domain was not attenuated. These results indicate that D614G adopts conformations that make virion membrane fusion with the target cell membrane more probable but that D614G retains susceptibility to therapies that disrupt interaction of the SARS-CoV-2 S protein with the ACE2 receptor.

## INTRODUCTION

Next-generation sequencing permits real-time detection of genetic variants that appear in pathogens during disease outbreaks. Tracking viral variants now constitutes a requisite component of the epidemiologist’s toolkit, one that can pinpoint the origin of a zoonotic virus and the trajectory it takes from one susceptible host to another (Hadfield et al., 2018; Shu and McCauley, 2017). Lagging behind sequence-based modeling of virus phylogenies and transmission chains is the ability to understand the effect of viral variants on the efficiency of transmission between hosts or on the clinical severity of infection. Most sequence variants that arise during virus replication are either detrimental to the fitness of the virus or without consequence. Even so, such variants can increase in frequency over the course of an outbreak by chance (Grubaugh et al., 2020). More rarely, though, increasing frequency of a variant can reflect competitive advantage due to higher intrinsic replication capacity, with increased viral load and transmissibility.

In December 2019, an outbreak of unexplained fatal pneumonia became apparent in Wuhan City, Hubei Province, China. By early January 2020, SARS-CoV-2 was identified as the virus causing the disease (Huang et al., 2020; Lu et al., 2020; Wu et al., 2020a, 2020b; Zhou et al., 2020b; Zhu et al., 2020). After SARS-CoV (Drosten et al., 2003; Ksiazek et al., 2003) and MERS-CoV (Zaki et al., 2012), SARS-CoV-2 is the third human coronavirus this century known to cause pneumonia with a significant case-fatality rate (Coronaviridae Study Group of the International Committee on Taxonomy of Viruses, 2020). Hundreds of coronaviruses have been identified in bats, including at least 50 SARS-like *Sarbecoviruses (Lu et al., 2020; Zhou et al., 2020a)*. The virus closest in sequence to SARS-CoV-2 observed to date was isolated from a bat (Zhou et al., 2020b) though the most proximal animal reservoir for SARS-CoV-2 remains unknown (Andersen et al., 2020; Lam et al., 2020).

Over the course of the SARS-CoV-2 pandemic, there have been reports of super-spreaders and transmission chains that have been more difficult to interrupt in some locations than in others (Hu et al., 2020). As few as 10% of SARS-CoV-2 infected people account for the majority of virus transmission events (Endo et al., 2020). Reported case fatality rates have varied by more than 10-fold (Center for Systems Science and Engineering at Johns Hopkins University; COVID-19 National Emergency Response Center, Epidemiology and Case Management Team, Korea Centers for Disease Control and Prevention, 2020; Grasselli et al., 2020; Onder et al., 2020). Such differences may reflect regional differences in the age of susceptible populations or the prevalence of comorbidities such as hypertension. Regional variation in diagnostic testing frequency, or in rate of reporting, is another likely explanation for these differences, since a wider net of diagnostic testing will capture milder cases (Zhao et al., 2020). Understanding the cause of these observed differences is critical for rational utilization of limited medical resources to counteract the pandemic.

Coronaviruses have the largest genomes of any known RNA viruses (Saberi et al., 2018) and they encode a 3’-to-5’-exoribonuclease required for high-fidelity replication by the viral RNA-dependent RNA polymerase (Denison et al., 2011; Smith et al., 2014). By preventing otherwise lethal mutagenesis (Smith et al., 2013), the repair function of the coronavirus exonuclease is thought necessary for the coronavirus genome size to extend beyond the theoretical limit imposed by error rates of viral RNA polymerases (Holmes, 2003). Though the rate of sequence variation among SARS-CoV-2 isolates is modest, over the course of the pandemic the virus has had opportunity to generate numerous sequence variants, many of which have been identified among the thousands of SARS-CoV-2 genomes posted for public access (https://www.gisaid.org/) (Hadfield et al., 2018). Here we investigate potential functional consequences of one of these variants, the Spike protein variant D614G.

## RESULTS

### The SARS-CoV-2 D614G S protein variant supplanted the ancestral virus

Minimal genetic variation among SARS-CoV-2 genomes sequenced from the earliest COVID-19 cases suggests that a recent common ancestor gave rise to all SARS-CoV-2 genomes seen during the outbreak (Rambaut et al., 2020). Over the course of the pandemic, a total of 12,379 distinct single nucleotide polymorphisms (SNPs) have been identified in genomic data (relative to the ancestral genome NC_045512.2; GISAID download on 25 June 2020), 99.5% of which are within viral genes, and 63.3% of which encode amino acid changes. 6,077 of the SNPs were seen only once in the dataset and only four SNPs rose to high frequency. Among the latter, A23403G was first reported at the end of January 2020 in virus genomes from China (hCoV-19/Zhejiang/HZ103/2020; 24 January 2020) and Germany (hCoV-19/Germany/BavPat1-ChVir929/2020; 28 January 2020). Given how few SARS-CoV-2 genomes have been sequenced from early in the outbreak, the geographic origin of A23403G cannot be determined.

The frequency of A23403G has increased steadily over time. It is now present in approximately 74% of all published sequences (Figure 1A). Sequences that have become available over recent weeks indicate that it has nearly reached fixation globally (Figure 1A), as well as separately within Europe, North America, Oceania, and Asia (Figure 1B). A23403G emerged at higher frequency in Africa and South America, suggesting a founder effect where a plurality of introductions carried it to these regions. Although recent data for South America are sparse, available sequences suggest A23403G is approaching fixation in this continent as well. Thus, A23403G is now universally common on every continent except Antarctica.

**Figure 1.**
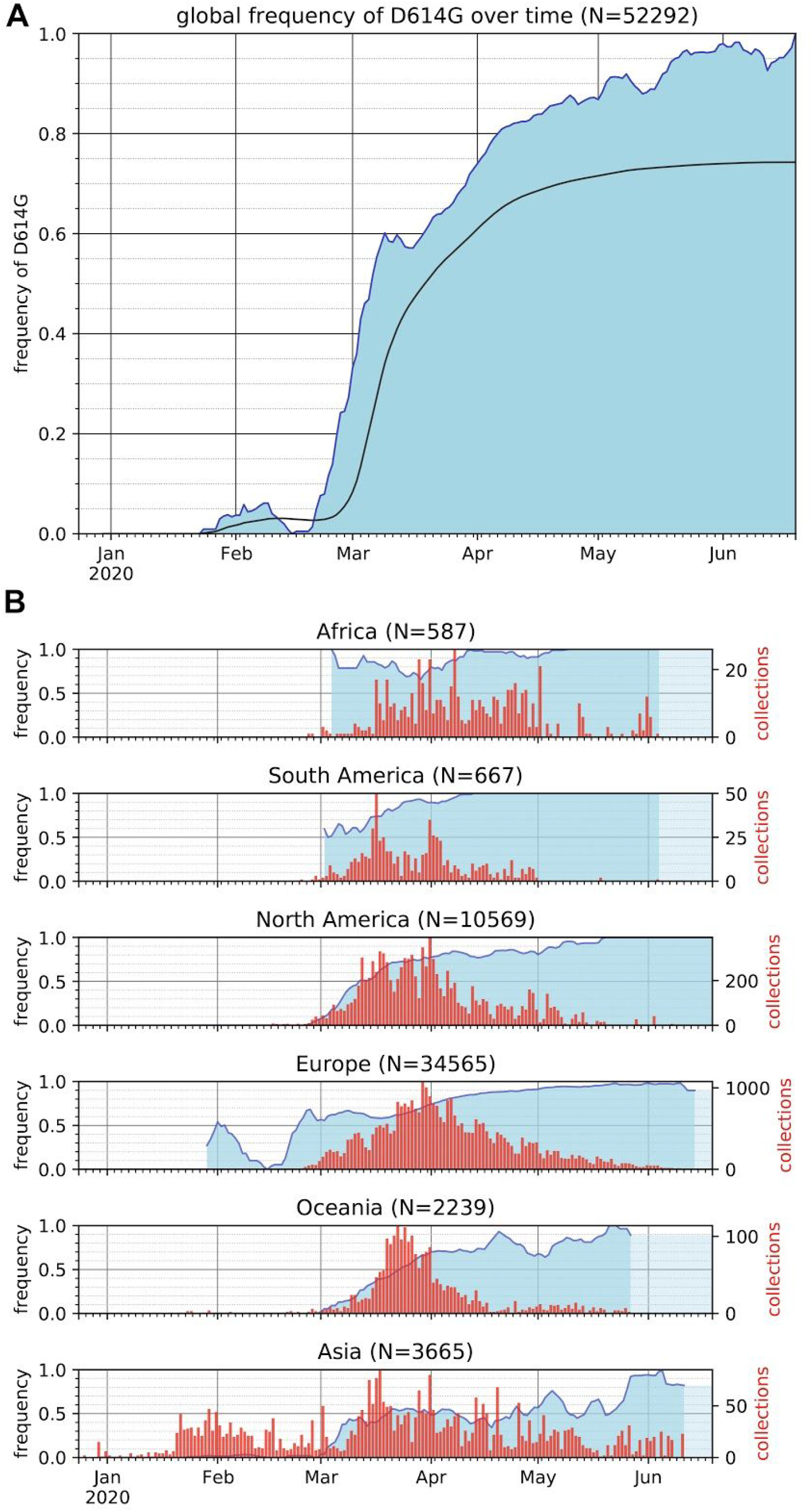
Over the course of the COVID-19 pandemic the frequency of the SARS-CoV-2 S protein D614G variant has risen to near fixation. (A) The global frequency of the S protein D614G variant over time in sequences published via the GISAID SARS-CoV-2 database, as of 25 June 2020. The filled plot (blue) represents a 7-day rolling average of the fraction of sequences bearing the D614G variant for each collection date. Dates without published sequences are linearly interpolated. The cumulative frequency of the D614G variant (black line) is overlaid, showing the frequency of D614G in sequences from samples collected up to and including each date. (B) The frequency of the D614G variant over time (blue) in sequences collected from six continental regions, using the same dataset as in (A), plotted as a 7-day rolling average. Frequencies are interpolated for dates without published sequence. The frequency of the last date with data is carried forward where recent dates lack data to indicate the most recent calculated frequency (light blue). The number of samples collected, sequenced, and published is overlaid (red bars) to provide the denominator used in calculating the frequency for each date, and to illustrate the non-uniformity of sampling.

### The SARS-CoV-2 D614G S protein variant increases infectivity in cell culture

The non-synonymous nucleotide change A23403G encodes the SARS-CoV-2 spike (S) protein variant D614G. The ability of the D614G S protein variant to target virions for infection of ACE2-positive cells was assessed using single-cycle lentiviral vector pseudotypes in tissue culture. Mammalian expression plasmids were engineered to encode the ancestral S protein D614 or the D614G variant. Each S protein expression plasmid was separately transfected into HEK-293 cells with plasmids encoding HIV-1 structural proteins and enzymes. Separate plasmids were transfected that encode RNAs with HIV-1 *cis*-acting signals for packaging and replication, and either GFP or luciferase (Luc) reporter cassettes. For each condition tested, multiple virus stocks were produced and tested individually after vector particle normalization using reverse transcriptase activity. 48 hours after challenge with the vectors, the transduction efficiency of each virion preparation was assessed by measuring the percent GFP-positive cells using flow cytometry, or by quantitating target cell-associated luciferase activity.

When Calu-3 human lung epithelial cells were used as targets, challenge with virus bearing D614G resulted in 6-fold more GFP-positive cells, or 5-fold more bulk luciferase activity, than did particles bearing D614 S protein (Figure 2A). When Caco-2 human colon epithelial cells were used as target cells, 4-fold higher infectivity was observed with D614G (Figure 2A). Additionally, when HEK-293 cells or SupT1 cells had been rendered infectable by stable expression of exogenous ACE2 and TMPRSS2, D614G was 9-fold more infectious than D614 (Figure 2A).

**Figure 2.**
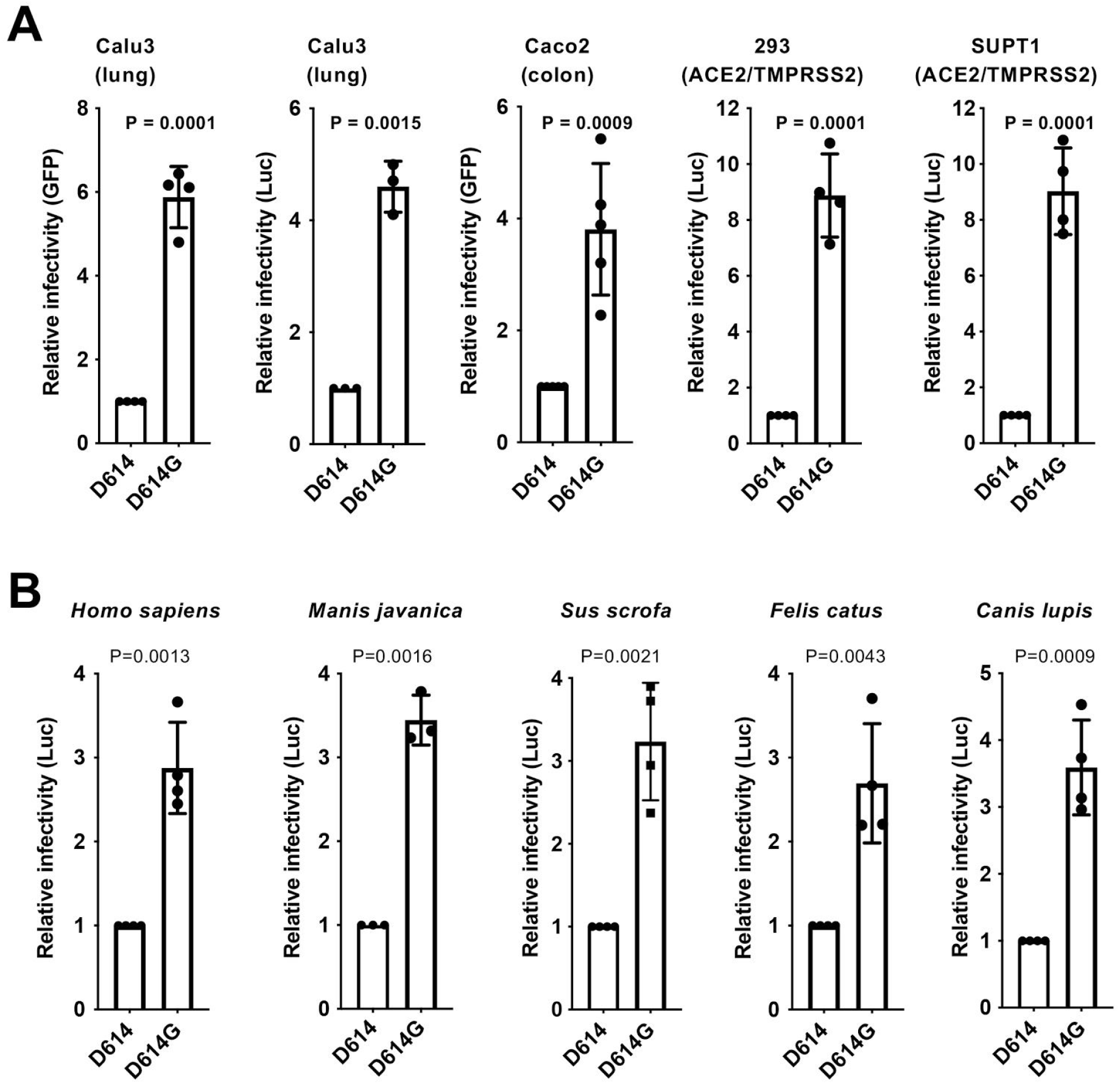
SARS-CoV-2 D614G S protein variant enhances infectivity. A. Lentiviral virions bearing either GFP or Luciferase transgenes, and pseudotyped with either SARS-CoV-2 D614 or D614G S proteins, were produced by transfection of HEK293 cells, and used to transduce human Calu3 lung cells, Caco2 colon cells, and either HEK293 or SupT1 cells stably expressing ACE2 and TMPRSS2. Relative infectivity of D614G vs D614, with D614 set at 1, was determined based on flow cytometry for percent GFP-positivity or bulk luciferase activity. Each point represents transduction with a lentiviral stock that was derived from an independent transfection. Shown are the mean +/- SD. B. Lentiviral virions bearing a luciferase transgene, pseudotyped with either SARS-CoV-2 D614 or D614G S proteins, were produced by transfection of HEK293 cells, and used to transduce human HEK293 cells transiently transfected with plasmids encoding the indicated ACE2 orthologs. Relative infectivity of D614G vs D614, with D614 set at 1, was determined based on bulk luciferase activity. Each point represents transduction using lentiviral stock derived from an independent transfection. Shown are the mean +/- SD.

### D614G increases infectivity on target cells bearing different ACE2 orthologs

The likely zoonotic origin of SARS-CoV-2 raises the question whether D614G was selected during the pandemic as a result of human-to-human transmission. Structural and biochemical studies (Letko et al., 2020; Walls et al., 2020; Wrapp et al., 2020; Zhou et al., 2020b) have shown that SARS-CoV-2 bears a receptor binding domain with high affinity for ACE2 from humans, ferrets, cats, and other species that encode similar ACE2 (Wan et al., 2020). In a heterologous expression system, human, civet, horse-shoe bat, and pig orthologs each conferred susceptibility to infection, while the mouse ortholog did not (Zhou et al., 2020b). To determine whether the increased infectivity of D614G is specific for certain ACE2 orthologs, HEK-293 were transfected separately with plasmids encoding ACE2 orthologs from human (*Homo sapiens*), Malayan pangolin (*Manis javanica*), pig (*Sus scrofa*), cat (*Felis catus*), dog (*Canis lupis*), rat (*Rattus norvegicus*), or mouse (*Mus musculus*). Each ACE2 transfectant was then challenged with luciferase reporter viruses bearing SARS-CoV-2 Spike protein, either D614 or D614G. Relative increase in infectivity due to D614G was comparable in cells expressing human, pangolin, pig, cat, or dog ACE2 orthologs (Figure 2B). The relative increase in infectivity due to D614G on cells bearing rat or mouse ACE2 orthologs could not be quantitated since, in both cases, the luciferase activity with D614 was too close to background. These results suggest that increased infectivity due to D614G is not specific for human ACE2.

### D614G does not alter S protein synthesis, processing, and incorporation into SARS-CoV-2 particles

The effect of D614G on S protein synthesis, processing, and incorporation into virion particles produced by SARS-CoV-2 structural proteins was assessed next (Figure 3A). HEK-293T cells were transfected with plasmids expressing each of the four SARS-CoV-2 virion-associated structural proteins (Figure 3A). Western blots were then performed on the cell lysate and on proteins from the pellet after ultracentrifugation of the transfection cell supernatant. The minimal requirement for virion assembly by other coronaviruses is the membrane protein (M) and the envelope protein (E) (for mouse hepatitis virus (Bos et al., 1996; Vennema et al., 1996)), or the M and nucleocapsid protein (N) (for SARS-CoV (Huang et al., 2004)). In the case of SARS-CoV-2, M protein production was sufficient to release particles from transfected cells, though particle release by M was increased by co-production of N (Figure 3B). The S proteins D614 and D614G were produced to comparable levels, processed to S1 and S2 with comparable efficiency, and incorporated into SARS-CoV-2 virus-like particles at similar levels (Figure 3B). These results suggest that increased infectivity due to D614G is primarily manifest after virion assembly, during entry into target cells.

**Figure 3.**
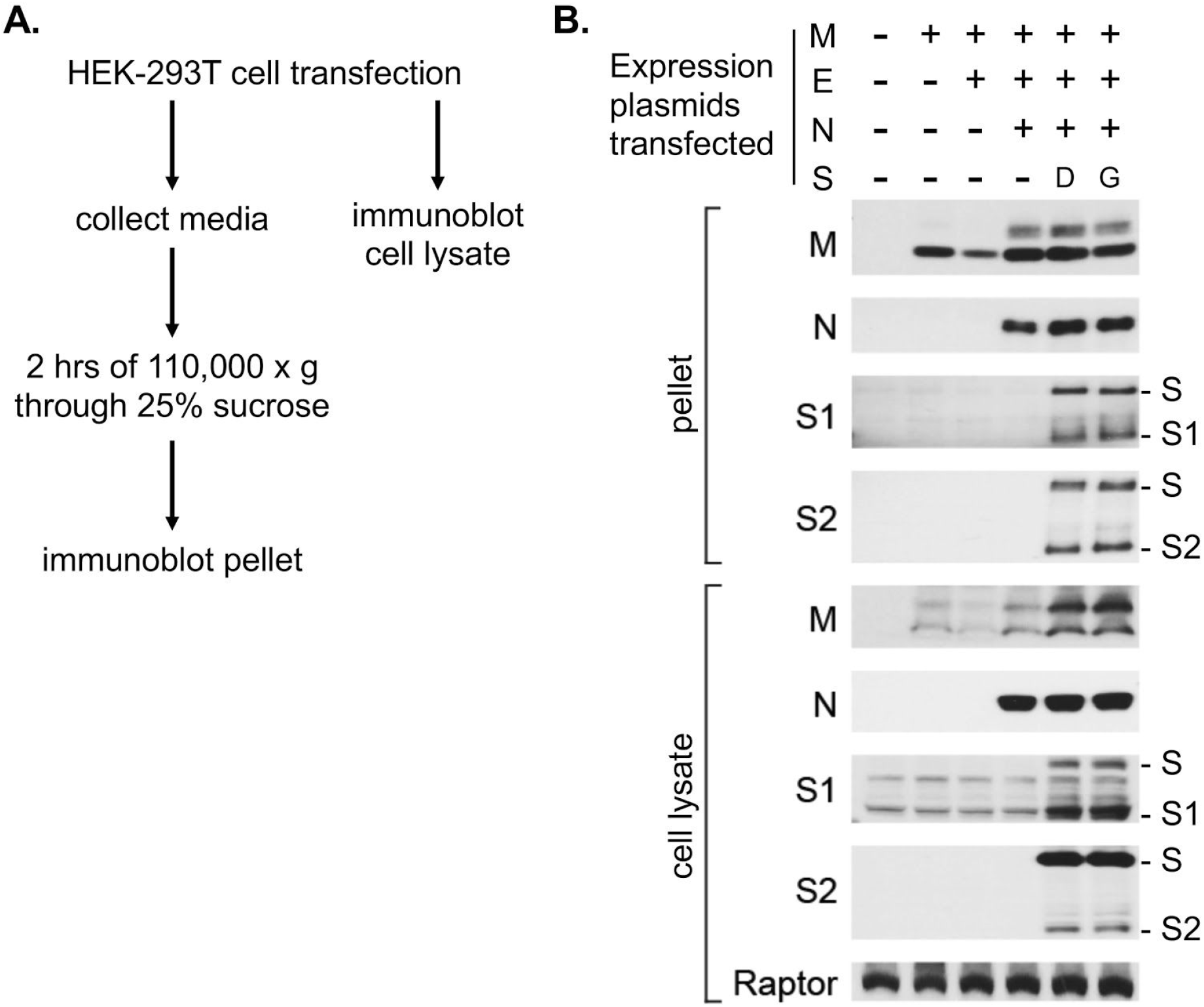
Synthesis, processing, and incorporation of S protein variants into SARS-CoV-2 virus-like particles. (A) Schematic showing how SARS-CoV-2 structural proteins were produced in HEK-293T cells and virus-like particles were enriched from the supernatant by ultracentrifugation. (B) HEK-293T cells were transfected with plasmids encoding the proteins indicated at the top. Western blots were then performed on cell lysate and on ultracentrifuge pellets from cell culture supernatant with the primary antibodies indicated on the left of the blots. Anti-Raptor antibody was used as a loading control for the cell lysate. Uncleaved S protein, as well as the S1 and S2 cleavage products, are indicated on the right.

### D614G decreases the affinity for ACE2 by increasing the rate of dissociation

The receptor-binding domain of the S protein interacts directly with ACE2, the primary entry receptor for SARS-CoV and SARS-CoV-2 on susceptible target cells (Hoffmann et al., 2020; Letko et al., 2020; Tian et al., 2020; Wrapp et al., 2020; Zhou et al., 2020b). Though D614G is located outside of the receptor-binding domain, this non-conservative amino acid change might alter ACE2-binding properties via allosteric effects. Surface plasmon resonance (SPR) was used to determine if the kinetics of SARS-CoV-2 S protein binding to human ACE2 is changed by D614G. Human ACE2 was immobilized and the binding of soluble, trimeric SARS-CoV-2 S protein, either D614 or D614G, was detected. At 25°C, the rate of association with ACE2 was little different between D614G and D614, but D614G dissociated from ACE2 at a rate four-fold faster than D614, resulting in a 5.7-fold reduction in binding affinity (Figure 4). At 37°C, the association rate between D614G and ACE2 was slower than between D614 and ACE2, and the dissociation rate of D614G was faster, again resulting in five-fold reduction in binding affinity (Figure 4). These data demonstrate that the increased infectivity of D614G is not explained by greater ACE2 binding strength.

**Figure 4.**
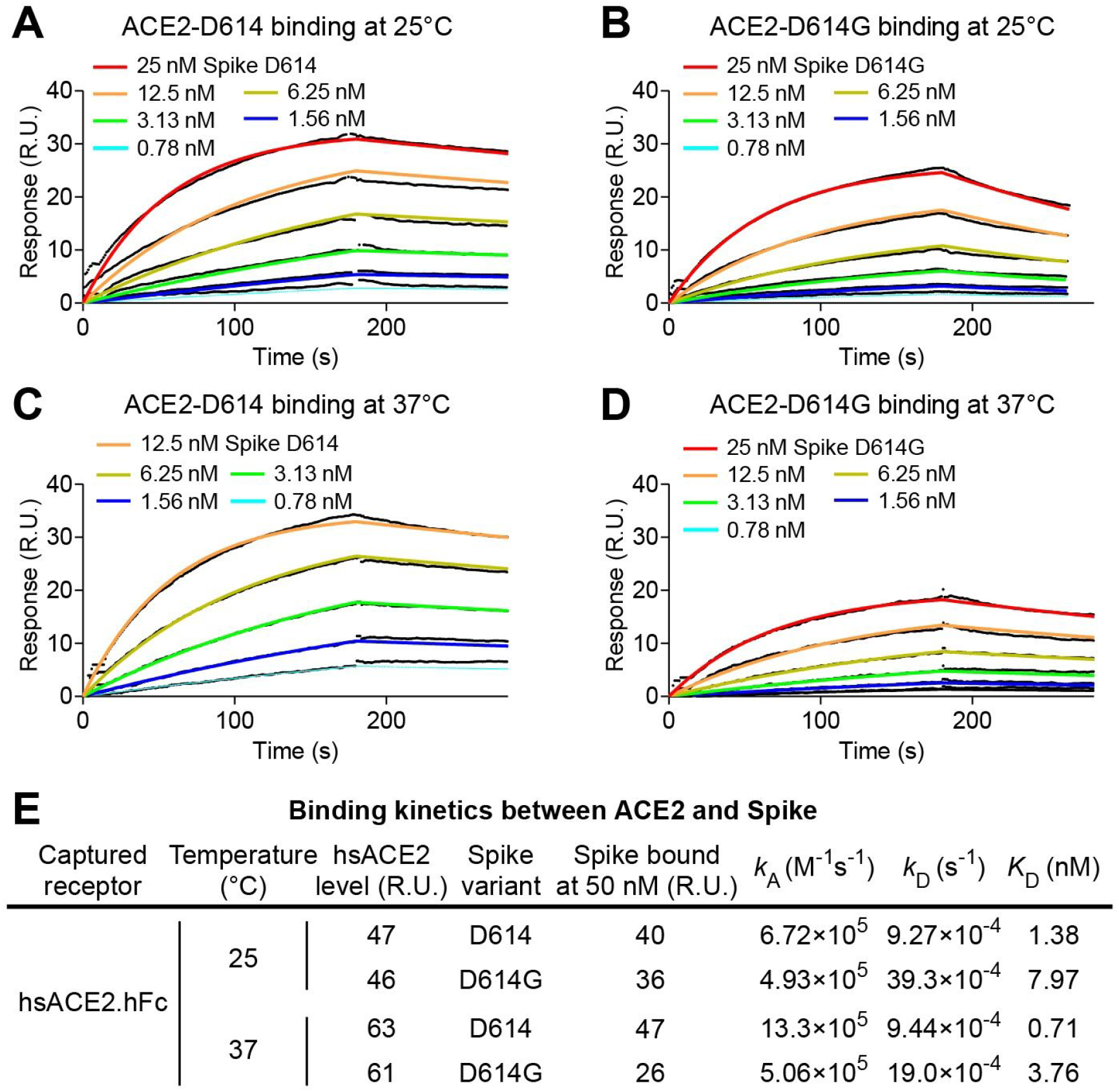
SARS-CoV-2 D614G S protein variant binds ACE2 weaker than the ancestral protein. **(A-D)** SPR measurement of D614-ACE2 binding (A and C) and D614G-ACE2 binding (B and D), at 25°C (A and B) or at 37°C (C and D). **(E)** Summary of kinetic parameters measured in A-D. D614G binds ACE2 five-fold weaker than D614 at both temperatures tested.

### D614G and the ancestral S protein are equally sensitive to neutralization by monoclonal antibodies targeting the receptor binding domain

The global spread and enhanced infectivity of the SARS-CoV-2 D614G variant raises the question of whether this change in S protein structure may compromise the effectiveness of antiviral therapies targeting this viral protein, especially if they were designed to target D614. To determine if this is the case, the neutralization potency of four monoclonal antibodies that target the SARS-CoV-2 Spike protein receptor-binding domain was assessed. These fully human monoclonal antibodies are currently under evaluation in clinical trials (NCT04425629, NCT04426695) as therapeutics for COVID-19 (Hansen et al., 2020). Each of these monoclonal antibodies, whether tested individually or in various combinations, demonstrated similar neutralization potency against D614G as they did against D614 (Figure 5).

**Figure 5.**
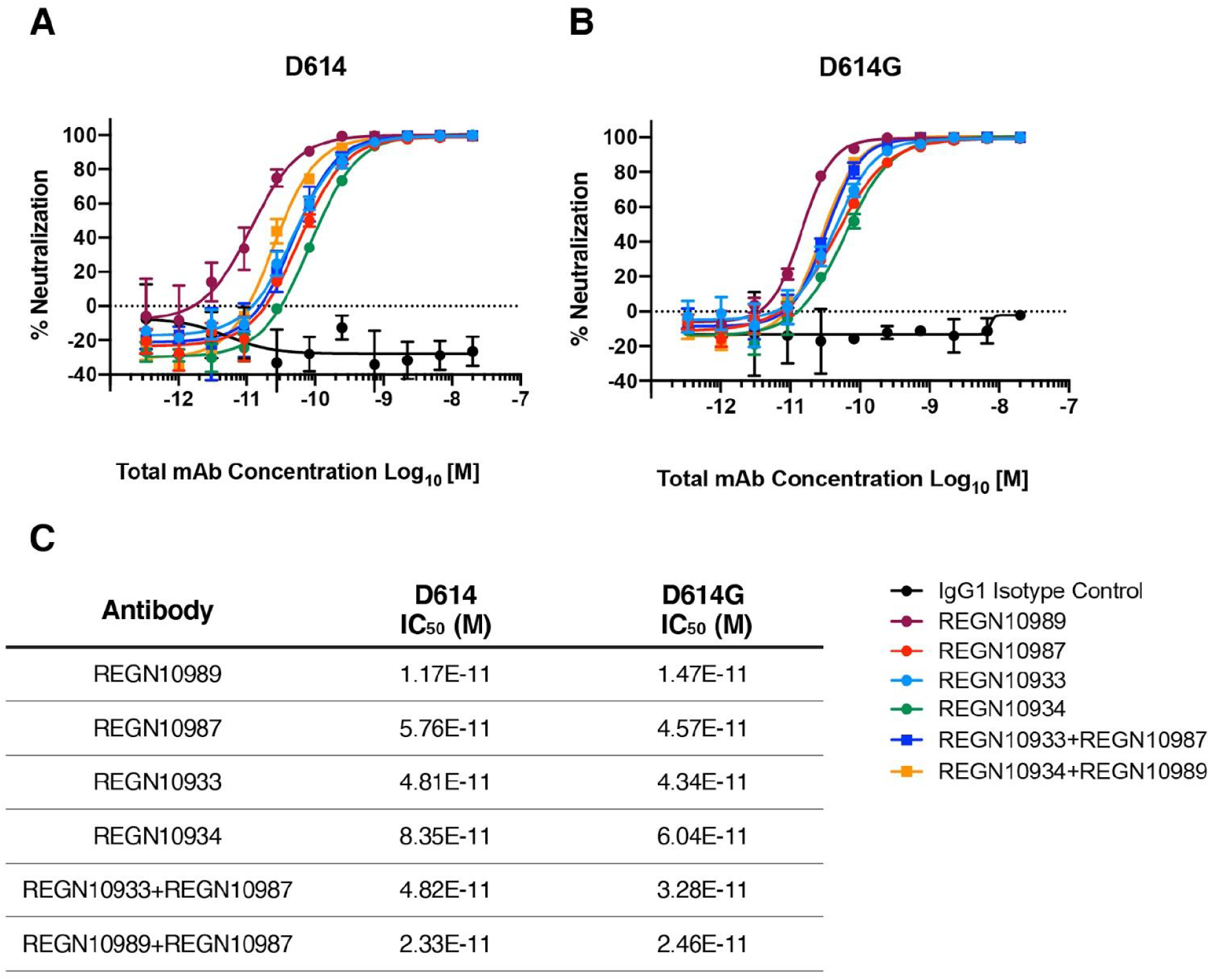
Neutralization potency of monoclonal antibodies targeting the SARS-CoV-2 S protein receptor binding domain. (A and B) Vero cells were challenged with pVSV-SARS-CoV-2-S-mNeon pseudoparticles encoding either D614 (A) or D614G S protein variants, in the presence of serial dilutions of the indicated human monoclonal antibodies targeting the SARS-CoV-2 S protein receptor binding domain, or IgG1 isotype control. mNeon protein fluorescence was measured 24 hours post-infection as a read-out for virus infectivity. Data is graphed as percent neutralization relative to virus only infection control. (C) Neutralization potency (IC_50_) of individual monoclonals and of combinations of monoclonals, against the SARS-CoV-2 D614G and D614G S protein variants, as indicated.

### D614G changes the conformation of the S1 domain in the SARS-CoV-2 S protein

Structural studies of the trimeric S protein ectodomain demonstrate that the receptor-binding domain of each protomer can independently adopt either a closed or an open conformation, giving rise to asymmetric trimers (Walls et al., 2020; Wrapp et al., 2020). The open conformation is believed to be required for ACE2-binding and to be on-pathway for S protein-mediated fusion of the virion membrane with the target cell membrane (Tortorici and Veesler, 2019). Since the increased infectivity of D614G was not explained by increased affinity for ACE2 (Figure 4), cryo-electron microscopy (cryo-EM) was used to illuminate potential structural features that distinguish D614G from D614. Both S protein variants, D614 and D614G, were expressed in mammalian cells as soluble trimers. When enriched from culture media, and eluted from a size exclusion column, single peaks were observed for each variant protein at ~500 kD, the predicted mass of the homotrimer (Figure 6A, blue line). Enrichment of the protein complexes, and integrity of the full-length protomers, was additionally confirmed by SDS-PAGE (Figure 6B).

**Figure 6.**
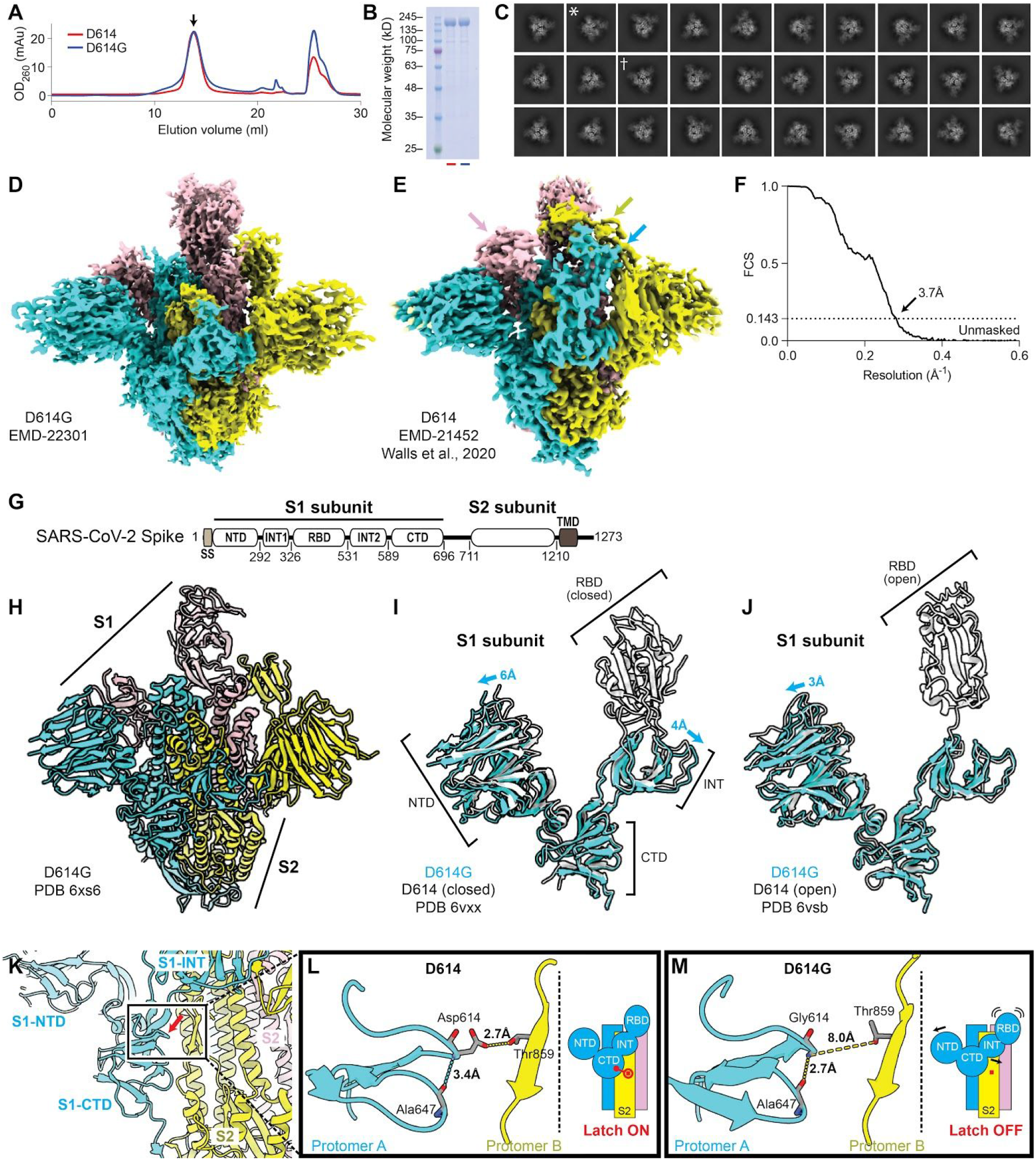
Structural determination of Spike D614G. (A) Size exclusion column elution profiles of D614G (blue line) and D614 (red line). (B) Coomassie-stained SDS-PAGE of the peaks collected in (A) shows equivalent, full-length monomeric S proteins at around 180 kD. (C) 2D-clustering of trimeric D614G particles. Dagger denotes a top-view. Asterisk denotes a side-view. (D) D614G envelope from the three-dimensional reconstruction. EMDB: EMD-22301. (E) Published D614 envelope (EMD-21452). Arrows point to the density corresponding to the receptor binding domains, which is missing in the corresponding positions in (D). (F) Gold-standard Fourier shell correlation of the density map in (D). (G) Domain arrangement of the SARS-CoV-2 S protein. (H) Atomic model for D614G without the receptor binding domain. PDB: 6XS6. (I-J) Comparison of the D614G S1 subunit with the closed conformation (I) and open conformation (J) of the D614 S1 subunit. Arrows indicate the relative movement of the S1 subunit of D614G. (K) Position of amino acid 614 on the S protein. (L-M) Substitution of Asp614 with glycine changes hydrogen bonding around residue 614. In the case of D614 (L), an inter-protomer hydrogen bond is detected. For D614G (M), the Asp614-Thr859 hydrogen bond is eliminated, and interaction with intradomain Ala647 is strengthened.

Well-defined particles of S protein trimers were evident by cryo-EM, and reference-free, two-dimensional clustering revealed structural details from different orientations (Figure 6C). Three-dimensional clustering and refinement generated the final density map for D614G (Figure 6D, EMD-22301), which showed a similar overall architecture to the published map of D614 (Figure 6E), but with distinct features, as discussed below. Unmasked Fourier shell correlation (FSC) analysis indicated that the D614G map had a mean resolution of 3.7 Å (gold-standard criteria, Figure 6F), which is sufficient to reveal fine differences from D614.

The SARS-CoV-2 S protein consists of S1 and S2 subunits, with multiple domains within S1 (Figure 6G). Comparing the map of D614G with that of D614, the N-terminal domain (NTD), the intermediary domain (INT), and the C-terminal domain (CTD) of the S1 subunit were clearly identified. However, the density corresponding to the receptor binding domain (RBD) that was well-resolved in D614 (arrows, Figure 6E), was missing in the D614G map, suggesting that the RBD is flexible and adopts multiple conformations (see section below).

Based on the resolved ensemble cryo-EM density map and the primary sequence, an atomic model of D614G without the RBD was built (Figure 6H, PDB: 6xs6). In the structural model, the S2 subunit of D614G overlapped well with the published structures for D614. In contrast, there was significant deviation within the S1 subunit (Figure 6I and 6J). When the S1 subunit of D614G was superimposed on the closed conformation of D614, the S1-NTD and S1-INT shifted away from each other by 6Å and 4Å, respectively (Figure 6I), revealing a wider space between these two domains. When the S1 subunit of D614G was superimposed on D614 in the open conformation, the S1-NTD was shifted outwards by 3Å, while the S1-INT overlapped well with that of D614 (Figure 6J). With respect to the ancestral protein, then, the D614G variant induces interdomain conformational changes, and likely leads to a more open conformation within the S1 subunit.

D614 localizes to the interface between two protomers where its side chain forms hydrogen bonds with the T859 side-chain in the adjacent protomer (Figure 6K). The effect of D614G on this interaction could be assessed since local resolution in the map generated here reaches 3.2Å. The atomic model showed that D614G has two consequences. First, D614G disrupts the inter-protomer hydrogen bond with Thr859 (Figure 6L and M), weakening the stability of the trimer. In effect, D614 acts as a “latch” that secures two protomers together, and D614G loosens this latch (Figure 6L and M). Second, the backbone amine of the glycine residue shortens the hydrogen bond with the carboxyl group of Ala647 within the same protomer, from 3.4 Å to 2.7 Å, and thus strengthens it. This interaction might internally stabilize the S1-CTD.

### D614G shifts the probability that SARS-CoV-2 S protein trimers occupy open conformations

The RBD cannot be visualized in the ensemble-averaged map for D614G. To better assess the conformation of the RBD, the flexible region of S1 was subjected to masked 3D classification and refinement, with the aim of resolving the conformational heterogeneity in that region. Two distinct classes arose from this analysis of the dataset. 58% of the protomers adopted an open conformation in which the RBD is positioned to interact with ACE2, and 42% were in the closed conformation in which the RBD is buried (Fig. 7A and 7B). This ratio of the two conformations contrasted dramatically with assessment of D614 structural data (Walls et al., 2020), in which only 18% of the RBDs were in the open conformation and 82% in the closed conformation. The closed conformation of D614G overlapped well with the closed conformation of D614, while the open conformation of D614G showed significant deviation of all S1 domains away from the S2 subunit when compared with the open conformation of D614 (Fig. 7C). This suggests that the D614-Thr859 hydrogen bond latches adjacent protomers together when one protomer is in the open conformation.

**Figure 7.**
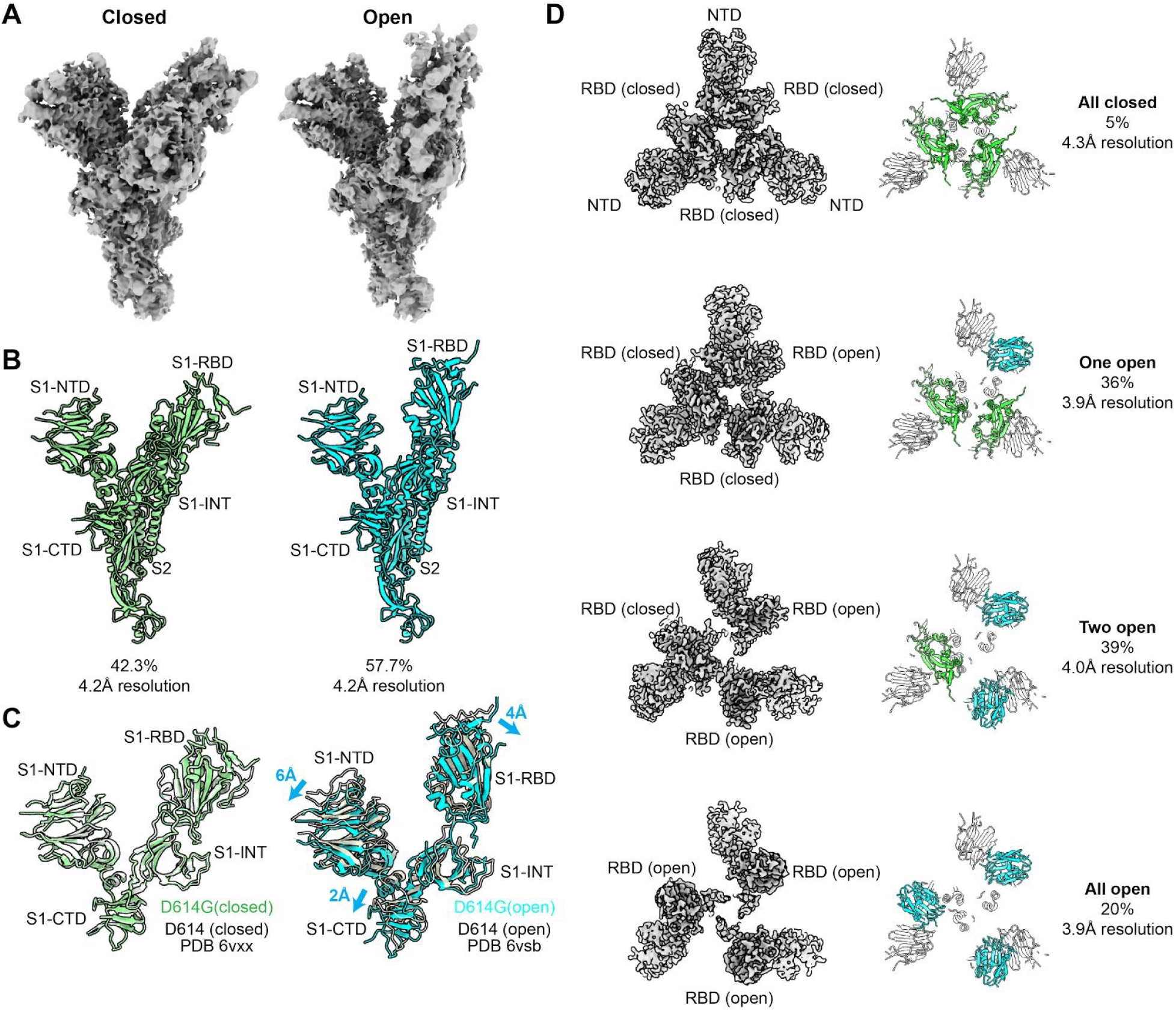
D614G populates more open conformations than does the ancestral S protein. (A) Cryo-EM density maps of the two conformations of D614G protomer. The first is a closed conformation with a buried RBD. The second is an open conformation with the RBD standing up. (B) Atomic models for the closed (left) and open (right) conformations for the two D614G protomers shown in (A). (C) Comparison of the two D614G protomer S1 subunit conformations with the corresponding conformations of the D614 protomer S1 subunit. (D) The D614G S protein trimer adopts four conformations. In addition to the all closed and one open conformation detected with the D614 S protein trimer, the D614G S protein trimer adopts two-open and three-open conformations.

Further classification of D614G particles to determine the distribution of RBD conformations within trimers revealed four different categories (Figure 7D). 5% of the particles had all three protomers in the closed conformation. 36% had one open protomer. 39% had two open protomers. 20% of the particles had all three protomers in the open conformation (Fig. 7D). The same analysis of the D614 dataset (Walls et al., 2020) showed that 46% of particles were in the all closed conformation and 54% in the one open conformation. These data further emphasize the contrast between D614G conformational space and that of D614, and suggest that the greater flexibility of the D614G RBD confers increased infectivity to D614G.

## DISCUSSION

The SARS-CoV-2 S protein variant D614G is one of only four SNPs, out of the more than 12,000 to be reported in GISAID, that has risen to high frequency (Figure 1). This suggests that D614G confers a replication advantage to SARS-CoV-2, such that it increases the likelihood of human-to-human transmission. Data in which the presence of D614G correlates with increased rates of transmission through human populations would support this hypothesis. To date, though, datasets that are powered sufficiently, and that are appropriately controlled for age and other important variables, have not been forthcoming. Prospective comparison of D614G transmission to that of D614 will be unlikely given that D614G has nearly gone to fixation world-wide (Figure 1), with only rare reports of D614 over recent weeks. It should be noted, however, that the roughly 53,000 high-quality SARS-CoV-2 genomic sequences available from GISAID as of 25 June 2020 are only a narrow snapshot of the pandemic, constituting approximately 0.5% of the 9.6 million cases confirmed as of that date (Dong et al., 2020). Additional sequencing of archived samples might provide opportunity to pinpoint the origin of D614G or to improve understanding of its trajectory.

Indirect evidence that D614G is more transmissible was provided here by experiments with pseudotyped viruses showing that D614G transduces 3 to 9-fold more efficiently than does the ancestral S protein (Figure 2A). This effect was seen with a range of cellular targets, including lung and colon epithelial cells, and required ACE2 on the target cell. The increased infectivity of D614G was evident whether or not TMPRSS2 was expressed on the target cells (Figure 2A). Efforts are underway to compare the replication efficiency of D614G with that of D614 in the context of the nearly 30,000 nucleotide SARS-CoV-2 genome. Such reverse genetic experiments, though, are technically difficult and potentially confounded by acquisition of unnatural, tissue culture-adapted mutations during genome rescue and expansion in transformed cell lines, as has occurred during similar assessments of Ebola virus variants (Marzi et al., 2018; Ruedas et al., 2017). Consistent with the increased ability of D614G to infect cells in tissue culture, several studies suggest that D614G is associated with increased viral load in people with COVID-19 (Korber et al., 2020; McNamara et al., 2020; Wagner et al.), though these studies quantitated SARS-CoV-2 RNA and did not measure infectious virus.

If SARS-CoV-2 Spike D614G is an adaptive variant that was selected for increased human-to-human transmission, following spillover from an animal reservoir, one might expect that increased infectivity would only be evident on cells bearing ACE2 orthologs similar to the human. In contrast to the primate-specific increase in infectivity that was reported for the major clade-forming Ebola virus glycoprotein variant from the 2013-2016 West African outbreak (Diehl et al., 2016; Urbanowicz et al., 2016), the increased infectivity of D614G was equally evident on cells bearing ACE2 orthologs from a range of mammalian species (Figure 2B). Though absolute infectivity differed based on the affinity of the RBD for the particular ACE2 ortholog, the relative increase in infectivity due to D614G was the same (Figure 2B). The fact that mouse and rat ACE2 orthologs were non-permissive for D614, but gave detectable infection with D614G, underlines this point. Examination of D614G infectivity using ACE2 orthologs encoded by other mammalian species may provide further insight into the structure and function of the D614G variant.

Insight into the mechanism by which D614G increases infectivity was gleaned from cryo-EM studies of the SARS-CoV-2 S protein trimer. D614G exhibited striking conformational changes within individual S protein protomers (Figure 6I and J, Figure 7C) and within the trimers (Figure 7D), all of which may be attributable to disruption of the interprotomer latch between D614 in S1 and T859 in S2 (Figure 7K-M). Weakening of the interprotomer contacts results in increased distance between the protomers and a dramatic flip in the ratio of open to closed S protein particles, from 82% closed and 18% open for D614, to 42% closed and 58% open for D614G. Perhaps the most remarkable conformational change was that, when the S protein trimers were examined, in addition to the all closed conformation and the one open conformation previously reported for D614 (Walls et al., 2020), a significant fraction of D614G trimers populated a two open conformation (39%) and an all open state (20%). Previous structural studies of SARS-CoV and MERS-CoV S proteins identified similar conformations, with only MERS-CoV S protein exhibiting an all open state like that reported here (Gui et al., 2017; Pallesen et al., 2017; Walls et al., 2019; Yuan et al., 2017). These studies suggested that only upon transition of an RBD to the open conformation is the protomer capable of binding the ACE2 receptor. Models of coronavirus S-mediated membrane fusion describe ACE2 binding to all three RBD domains in the open conformation as destabilizing the pre-fusion S trimer, leading to dissociation of S1 from S2 and promoting transition to the post-fusion conformation (Pallesen et al., 2017; Walls et al., 2019). Thus, the open conformation likely reflects an intermediate that is on-pathway to S-mediated membrane fusion. The well-populated all open conformation of D614G (Figure 7D) therefore suggests that this S protein variant increases infectivity by making virion membrane fusion with the target cell membrane more probable.

The D614G and D614 variants are equally sensitive to neutralization by four human monoclonal antibodies targeting the S protein RBD that are currently in clinical trials for people with COVID-19 (Figure 5). These results are in keeping with the fact that the location of D614G within the S protein is remote from the receptor binding domain and that D614G affinity for ACE2 is less than that of D614 (Figure 4). To date, no studies have demonstrated a worse clinical outcome for infection with D614G, and given the more open conformation of its receptor-binding domain, one might predict that D614G will be at least as immunogenic as the ancestral form.

## Methods

### Analysis of D614G frequency in published data

The frequency of the SARS-CoV-2 D614G S protein variant in published genomic data was examined using the full Nextstrain-curated set of sequences available from GISAID as of 25 June 2020 (Hadfield et al., 2018; Shu and McCauley, 2017). Sequences were aligned to the ancestral reference sequence (NCBI GenBank accession NC_045512.2) using mafft v7.464 (Katoh and Standley, 2013) with the “--keeplength” and “--addfragments” parameters, which preserve the coordinate space of the reference sequence. To remove lower-quality sequences from the dataset, all sequences in the alignment were masked with ambiguous bases (‘N’) in the regions spanning the first 100bp and the last 50bp, as well as at error-prone sites located at the (1-indexed, NC_045512.2 coordinate space) positions 13402, 24389, 24390. Sequences shorter than 28kb or with >2% ambiguous bases were removed from the alignment. The frequency of D614G was calculated in the resulting data by extracting the sequence region corresponding to the gene for the S protein, spanning 21563-25384bp. These sequences were processed using a script importing biopython (Cock et al., 2009) to remove any gaps introduced by the alignment process and translate the sequence to protein space. The identity of the variant at amino acid position 614 was tabulated for the full dataset and reported as frequency by date using the collection dates reported in the Nextstrain-curated metadata file available from GISAID (Hadfield et al., 2018; Shu and McCauley, 2017). The frequency was calculated as (# sequences with D614G)/(# sequences). Frequency within the six continental regions was calculated based on the “region” geographic classification associated with each sample in the metadata. Frequency values were linearly interpolated for dates surrounded by valid data. The frequency of the last date with data was carried forward where recent dates lack data. The bresulting values were rendered as plots using matplotlib (Hunter, 2007). The script for analyzing and plotting D614G variant frequency is available via GitHub: https://github.com/broadinstitute/sc2-variation-scripts

The diversity of SNPs and their functional effects based on the same GISAID sequences and MAFFT alignment used to plot the frequency of D614G over time, with the 5’ and 3’ ends not masked. In the alignment, ambiguous nucleotide codes (R,Y,W,S,M,K,H,B,V,D) were all masked with “N” values. SNPs were calculated from the alignment using the *snp-sites* tool (Page et al., 2016). The resulting VCF-format file was normalized using *bcftools* (Li et al., 2009) to include only SNPs. The VCF file with SNPs was annotated for functional effects using *SnpEff* (Cingolani et al., 2012).

### Plasmids

The plasmids used here were either previously described or were generated using standard cloning methods. The full list of plasmids used here is provided in the Key Resources Table. All newly engineered plasmids are available from public repositories, along with full sequences, at either https://www.addgene.org/Jeremy_Luban/ or https://www.aidsreagent.org (search under Jeremy Luban).

### Cell culture

All cells were cultured in humidified incubators with 5% CO_2_ at 37° C, and monitored for mycoplasma contamination using the Mycoplasma Detection kit (Lonza LT07-318). HEK293 cells (ATCC CRL-1573), and HEK-293T cells (CRL-3216 or CRL-11268) were cultured in DMEM supplemented with 10% heat-inactivated FBS, 1 mM sodium pyruvate, 20 mM GlutaMAX, 1× MEM non-essential amino acids, and 25 mM HEPES, pH 7.2. Calu3 cells (ATCC HTB-55) were maintained in EMEM supplemented with 10% FBS. Caco2 cells (ATCC HTB-37) were maintained in EMEM supplemented with 20% FBS. SUP-T1 [VB] cells (ATCC CRL-1942) were cultured in RPMI supplemented with 10% heat-inactivated FBS, 1mM sodium pyruvate, 20mM GlutaMAX, 1xMEM non-essential amino acids, and 25 mM HEPES, pH7.2. Vero cells (ATCC CCL-81) were cultured in DMEM high glucose media containing 10% heat-inactivated fetal bovine serum, and 1X Penicillin/Streptomycin/L-Glutamine.

### Virus production

24 hrs prior to transfection, 6 × 10^5^ HEK-293 cells were plated per well in 6 well plates. All transfections used 2.49 μg plasmid DNA with 6.25 μL TransIT LT1 transfection reagent (Mirus, Madison, WI) in 250 μL Opti-MEM (Gibco).

Single-cycle HIV-1 vectors pseudotyped with SARS-CoV-2 Spike protein, either D614 or D614G, were produced by transfection of either HIV-1 pNL4-3 Δenv Δvpr luciferase reporter plasmid (pNL4-3.Luc.R-E-), or pUC57mini NL4-3 Δenv eGFP reporter plasmid, in combination with the indicated Spike expression plasmid, at a ratio of 4:1.

ACE2 expression vectors were produced by transfecting cells with one of the pscALPSpuro-ACE2 plasmids, along with the HIV-1 *gag-pol* expression plasmid psPAX2, and the VSV glycoprotein expression plasmid pMD2.G (4:3:1 ratio of plasmids).. 16 hrs post-transfection, culture media was changed. Viral supernatant was harvested 48 hours after media change, passed through a 0.45 μm filter, and stored at 4°C. TMPRSS2 expression transfer vector was produced similarly but with pscALPSblasti-TMPRSS2.

### Exogenous reverse transcriptase assay

5 μL transfection supernatant was mixed with 5 μL 0.25% Triton X-100, 50 mM KCl, 100 mM Tris-HCl pH 7.4, and 0.4 U/μL RiboLock RNase inhibitor, and then diluted 1:100 in 5 mM (NH_4_)_2_SO_4_, 20 mM KCl, and 20 mM Tris-HCl pH 8.3. 10 μL of this was then added to a single-step, RT-PCR assay with 35 nM MS2 RNA (IDT) as template, 500 nM of each primer (5’-TCCTGCTCAACTTCCTGTCGAG-3’ and 5’-CACAGGTCAAACCTCCTAGGAATG-3’), and 0.1 μL hot-start Taq DNA polymerase (Promega, Madison, WI) in 20 mM Tris-Cl pH 8.3, 5 mM (NH_4_)_2_SO_4_, 20 mM KCl, 5 mM MgCl_2_, 0.1 mg/ml BSA, 1/20,000 SYBR Green I (Invitrogen), and 200 μM dNTPs in total 20 μL reaction. The RT-PCR reaction was carried out in a Biorad CFX96 real-time PCR detection system with the following parameters: 42°C for 20 min, 95°C for 2 min, and 40 cycles [95°C for 5 sec, 60°C for 5 sec, 72°C for 15 sec, and acquisition at 80°C for 5 sec].

### Generation of cell lines expressing ACE2 and TMPRSS2

2.5 x 10^5^ HEK-293 cells were plated per well in a 12 well plate. The next day cells were transduced with 250 uL of supernatant containing TMPRSS2-encoding lentivirus for 16 hr at 37°C, after which fresh media was added to cells. 48 hrs after transduction cells were replated and selected with blasticidin (InvivoGen, catalogue #ant-bl-1) at 10 ug/ml. After selection, cells were transduced similarly with supernatant containing ACE2-encoding lentivirus and selected with 1 ug/mL of puromycin (InvivoGen, San Diego, CA, catalogue #ant-pr-1).

1 x 10^6^ SupT1 cells were transduced with 400 uL of supernatant containing TMPRSS2-encoding virus followed by selection with 10 ug/mL blasticidin 48 hrs later. TMPRSS2 expressing SupT1 cells were then transduced with a second vector expressing ACE2, followed by puromycin selection at 1 ug/mL.

### Virus Infectivity Assays

16 hours prior to transduction, adherent cells were seeded in 96 well plates. HEK-293 cells were plated at 5 x 10^4^ cells per well. Calu3 and Caco2 were plated at 3 x 10^4^ per well. Cells were incubated in virus-containing media for 16 hrs at 37°C when fresh medium was added to cells. 48 to 72 hours after transduction cells were assessed for luciferase activity or for GFP by flow cytometry. For transfection of SupT1 cells, 1 x 10^5^ cells were plated in a U-bottom 96 well plate and spinfected at 30°C with 100 uL of virus-containing media for 2 hours at 1,200 x g. Fresh media was added after spinfection and cells were incubated 48-72 hours prior to analysis. Cells transduced with GFP virus were fixed with BD Cytofix (BD Biosciences, San Jose, CA, Cat number 554655) and analyzed using the Accuri C6system. Data was analyzed using FlowJo 10.5 (FlowJo, LLC, Ashland, OR). Cells transduced with luciferase expressing virus were assessed using Promega Steady-Glo system (Promega Madison, WI)

### Neutralizations assays with human monoclonal antibodies targeting the SARS-CoV-2 S protein receptor binding domain

VSV-SARS-CoV-2-S pseudoparticle generation and neutralization assays were performed as previously described (Baum et al., 2020; Hansen et al., 2020). HEK293T cells were seeded overnight in DMEM high glucose media (Life Technologies) containing 10% heat-inactivated fetal bovine serum (Life Technologies), and Penicillin/-Streptomycin-L-Glutamine (Life Technologies). The following day, Spike expression plasmids were transfected with Lipofectamine LTX (Life Technologies) following the manufacturer’s protocol. At 24 hours post transfection, the cells were washed with phosphate buffered saline (PBS) and infected at an MOI of 1 with the VSVΔG:mNeon/VSV-G virus diluted in 10 mL Opti-MEM (Life Technologies). The cells were incubated 1 hour at 37°C with 5% CO2. Cells were washed three times with PBS to remove residual input virus and overlaid with DMEM high glucose media (Life Technologies) with 0.7% low IgG BSA (Sigma), sodium pyruvate (Life Technologies), and gentamicin (Life Technologies). After 24 hours at 37° C with 5% CO2, the supernatant containing pseudoparticles was collected, centrifuged at 3,000 x g for 5 minutes to clarify, aliquoted, and frozen at −80° C.

For neutralization assays, Vero cells were seeded in 96-well plates 24 hours prior to assay and grown to 85% confluence before challenge. Antibodies were diluted in DMEM high 3 glucose media containing 0.7% Low IgG BSA (Sigma), 1X Sodium Pyruvate, and 0.5% Gentamicin (this will be referred to as “Infection Media”) to 2X assay concentration and diluted 3-fold down in Infection Media, for an 11-point dilution curve in the assay beginning at 10 ug/mL (66.67 nM). pVSV-SARS-CoV-2-S pseudoparticles were diluted 1:1 in Infection Media for a fluorescent focus (ffu) count in the assay of ~1000 ffu. Antibody dilutions were mixed 1:1 with pseudoparticles for 30 minutes at room temperature prior to addition onto Vero cells. Cells were incubated at 37° C, 5% CO2 for 24 hours. Supernatant was removed from cells and replaced with 100 uL PBS, and fluorescent foci were quantitated using the SpectraMax i3 plate reader with MiniMax imaging cytometer.

### Production of SARS-CoV-2 virus-like particles (VLPs)

HEK-293T cells were cultured in DMEM supplemented with 10% heat-inactivated bovine serum, and transfected with pcDNA3.1 plasmids encoding the SARS-CoV-2 M, E, N, and S proteins, in different combinations, as indicated. 1 ug of each plasmid was used, with 5 ug of total plasmid in each transfection, normalized using empty vectors, in 400 ul Opti-MEM and 18 ul of PEI. The transfection mixture was incubated at room temperature for 15 minutes and dropped into a 50% confluent 10 cm plate of HEK-293T cells. The media and cell lysate were collected after 60 hours. 10 ml of supernatant was passed through a 0.45 um syringe filter, layered on top of a 20% sucrose cushion in PBS, and spun at 30,000 rpm in an SW41 rotor (111,000 x g, average) for two hours. The pellet was washed once with ice cold PBS, resuspended in 150 ul of 2x SDS loading buffer (Morris formulation), and sonicated in an ice-water bath (Branson 2800) for 15 minutes. After removal of the supernatant, the transfected cells were lysed in 800 ul PBS with 1% Triton and protease inhibitor (Roche cOmplete EDTA-free tablets), and cleared of debris by spinning in a table-top centrifuge (Eppendorf 5424R) at top speed (20,000 x g, average) for 10 minutes.

### Immunoblots of SARS-CoV-2 VLPs

Proteins in VLP and cell lysate samples were separated by SDS-PAGE, as follows: 20 ul of unboiled VLP and cell lysate samples on a 10-20% Tris-Glycine gel to probe for the M protein; 2 ul of boiled cell lysate and 5 ul of boiled VLP samples on a 12% Tris-Glycine gel to probe for the N protein; 20 ul of boiled lysate and VLP samples on a 10% Tris-Glycine gel to probe for the S protein; 5 ul of boiled lysate on a 10% Tris-Glycine gel to probe for Raptor, as a loading control. Proteins were electro-transferred from the gels to PVDF membrane, which was blocked with 5% milk in Tris-Buffered Saline, pH 8.0, with 0.1% Tween-20, and detected with the indicated antibodies.

### Surface plasmon resonance analysis

Binding kinetics and affinities for ACE2.Fc were assessed using surface plasmon resonance technology on a Biacore T200 instrument (GE Healthcare, Marlborough, MA) using a Series S CM5 sensor chip in filtered and degassed HBS-EP running buffer (10 mM HEPES, 150 mM NaCl, 3mM EDTA, 0.05% (v/v) polysorbate 20, pH 7.4). Capture sensor surfaces were prepared by covalently immobilizing with a mouse anti-human Fc mAb (REGN2567) on the chip surface using the standard amine coupling chemistry, reported previously (Johnsson et al., 1991). Following surface activation, the remaining active carboxyl groups on the CM5 chip surface were blocked by injecting 1 M ethanolamine, pH 8.0 for 7 minutes. A typical resonance unit (RU) signal of ~12,000 RU was achieved after the immobilization procedure. At the end of each cycle, the anti-human Fc surface was regenerated using a 12 second injection of 20 mM phosphoric acid. Following the capture of the ACE2.Fc on the anti-human Fc mAb immobilized surface, 0.78 nM - 50 nM, two-fold serial dilutions, in duplicate, of soluble SARS-CoV-2 spike trimer protein, D614 or D614G, were injected for 3 minutes at a flow rate of 50 mL/min, with a 2 minute dissociation phase in the running buffer. All specific SPR binding sensorgrams were double-reference subtracted as reported previously (Myszka, 1999) and the kinetic parameters were obtained by globally fitting the double-reference subtracted data to a 1:1 binding model with mass transport limitation using Biacore T200 Evaluation software v 3.1 (GE Healthcare). The dissociation rate constant (*k_d_*) was determined by fitting the change in the binding response during the dissociation phase and the association rate constant (*k_a_*) was determined by globally fitting analyte binding at different concentrations. The equilibrium dissociation constant (*K_D_*) was calculated from the ratio of the *k_d_* and *k_a_*. The dissociative half-life (t½) in minutes was calculated as ln2/(*k_d_*\*60). The steady state analysis was performed using Scrubber software and the *K*_D_ value was determined.

### Production and enrichment of soluble SARS-CoV-2 S Protein trimers

FreeStyle 293-F cells were cultured in SMM-293 TII serum-free media (SinoBiological) and maintained in a 37°C shaker with 8% CO_2_ and 80% humidity. 600 ug of plasmid encoding His-tagged SARS-CoV-2 S protein was transfected into 400 ml of 293 FreeStyle cells at 10^6^ cells/ml, using 25 ml Opti-MEM and 1.8 ml PEI. 60 hours later, the media was collected and applied to 3 ml of Ni-NTA resin (Qiagen). The resin was collected and washed with PBS supplemented with 20 mM imidazole. Soluble S protein was eluted in PBS supplemented with 200 mM imidazole, and further enriched using a Superose 6 gel-filtration column (GE healthcare), in a buffer containing 25 mM NaHEPES (pH 7.4) and 150 mM NaCl. Yield was roughly 600 ug of soluble S protein homotrimer per liter of culture.

### CryoEM sample preparation and data collection

3 μL of enriched D614G trimers, at 2.5 mg/mL, was deposited on an UltrAuFoil R1.2/1.3 300 mesh grid that had been glow discharged for 30 seconds in a GloQube Plus Glow Discharge System. Plunge freezing was performed with a Vitrobot Mark IV using a blot force 0 and 5s blot time at 100% humidity and 4°C. Frozen grids were imaged with a Thermo Scientific Krios transmission electron microscope operated at 300 kV. Movies were acquired using EPU 2 software. 10,264 micrographs were automatically collected with a defocus ranging between 0.6 and 1.8 μm, at a nominal magnification of 155,000x with a pixel size of 0.52Å. 7,653 micrographs were collected with a tilt-angle of 30° to overcome the preferred orientation. The dose rate was set at 20.3 electron/Å^2^/s and total exposure time was 1.81s. With a dose fraction set at 62 frames, each movie series contained 62 frames and each frame received a dose of 0.6 electron/Å^2^.

### CryoEM data processing

Movie frame alignment, estimation of microscope contrast-transfer function parameters, particle picking, 2D classification, and homogeneous refinement using the published structure EMD-21452 as initial model were carried out in cryoSPARC. Ensemble averaging of 266,356 particles resulted in a 3.7Å map with C3 symmetry imposed according to “gold standard” Fourier shell correlation of 0.143. Local resolution was estimated using cryoSPARC to extend from 3Å to 6 Å. Particle stacks with well-refined orientation parameters were imported in Relion3.1. Focused 3D classifications with a soft mask on the S1 subunit of the protomer were performed on the C3 symmetry expanded particles. Two monomer conformations, namely the closed state (a) and the open state (b) were identified after 3 rounds of classification. Four trimer classes were identified by 3D classification on the trimer particles: class_3a (13,555 particles), class_3b (51,600 particles), class_2a1b (96,029 particles), class_1a2b (105,118 particles). Homogeneous refinements were then performed on these 4 classes, with EMD-21452 as the initial model in Relion3.1. C3 symmetry was imposed on the refinement of particles from class_3a and class_3b; C1 symmetry was imposed on the refinement of particles from class_2a1b and class_1a2b. The final resolutions were class_1a2b with 4Å, class_2a1b with 3.9Å, class_3a with 4.2Å and class_3b with 3.9Å. The final map was deposited with the accession code EMD-22301.

### Model building

Atomic models were prepared with Coot based on the resolved structure of D614 SARS-CoV-2 Spike (PDB: 6vxx and 6vsb). Real-space refinements were performed using PHENIX with secondary structure restraints. MolProbity was used to evaluate the geometries of the structural model. Corrected Fourier shell correlation curves were calculated using the refined atomic model and the cryo-EM density map. The coordinates were deposited with the accession code 6xs6.

## ACKNOWLEDGEMENTS

We thank all the members of the Sabeti, Shen, and Luban labs, and Thermo Fisher Scientific Workflow solutions team for technical assistance and helpful discussions. This work was supported by NIH grants R37AI147868 and R01AI148784 to J.L., NIH/NIAID U19AI110818 to P.C.S., 7DP2AI124384-02 to J.B.M., NCI K22CA241362 to K.S., a grant from the Evergrande COVID-19 Response Fund Award from the Massachusetts Consortium on Pathogen Readiness to J.L., a grant from the Worcester Foundation to K.S., a Sara Elizabeth O’Brien Fellowship Award/King Trust to L.Y., and a National Science Foundation Graduate Research Fellowship (Grant No. 1745303) to C.T.-T. A portion of this project has been funded in whole or in part with Federal funds from the Department of Health and Human Services; Office of the Assistant Secretary for Preparedness and Response; Biomedical Advanced Research and Development Authority, under OT number: HHSO100201700020C.

## AUTHOR CONTRIBUTIONS

P.S., C.A.K., K.S., and J.L. conceived the experiments. C.T.-T., S.F.S, J.E.L., and P.C.S. analyzed the SARS-CoV-2 sequence data. L.Y., T.N., Y.W., W.E.D., A.D., C.A.K., K.S., and J.L. designed and cloned the expression plasmids. L.Y., C.C., A.D., K.E.P., and A.B. performed the virology experiments. L.Y., A.D., and K.S. performed the virus biochemistry. A.R. performed the SPR experiments. K.V., S.B.E., and K.S. purified the proteins. X.W., A.B., and N.D. collected and processed cryo-EM data. S.B.E. and K.S. built the structural model. L.Y., C.H.T.-T., S.F.S, C.A.K., A.B., A.R., J.B.M., K.S., and J.L. wrote and edited the manuscript with input from all authors.

## DECLARATION OF INTERESTS

P.C.S. is a co-founder and shareholder of Sherlock Biosciences, and a Board member and shareholder of Danaher Corporation. J.E.L. consulted for Sherlock Biosciences. C.A.K., K.E.P., and A.B. are employed by Regeneron Pharmaceuticals and own stock options in the company. C.A.K. is an officer at Regeneron. X.W., A.B., and N.D. are employees of Thermo Fisher Scientific.

---

**Table 1.** Summary of cryo-EM data collection, 3D reconstruction, and model refinement

|  | **SARS-Covid19 Spike (D614G)** |
| --- | --- |
| **Imaging parameters and 3D reconstruction** |  |
| Cryo-EM grids | UltrAuFoil R1.2/1.3 |
| Calibrated magnification | 155000 |
| Acceleration voltage [kV] | 300 |
| Pixel size [Å] | 0.52 |
| Total dose [e<sup>-</sup>/Å<sup>2</sup>] | 37 |
| Exposure time [s] | 1.81 |
| Defocus range [μm] | -0.6 ~ -1.8 |
| Particles in final 3D reconstruction | 266356 |
| Resolution ('Gold-standard' at FSC 0.143) [Å] | 3.7 |
| **Model refinement** |  |
| Resolution in phenix.real_space_refine [Å] | 4.0 |
| No. atoms: protein | 18294 |
| Ligands / ions | N/A |
| r.m.s. deviations: bond lengths [Å] | 0.004 |
| r.m.s. deviations: bond angles [°] | 0.808 |
| MolProbity score | 4.49 |
| Rotamer outliers [%] | 0.00 |
| Ramachandran angles: favored [%] | 93.88 |
| Ramachandran angles: allowed [%] | 6.12 |
| Ramachandran angles: outliers [%] | 0.00 |
| **PDB** | 6xs6 |
| **EMDB** | EMD-22301 |

**Table 2.**
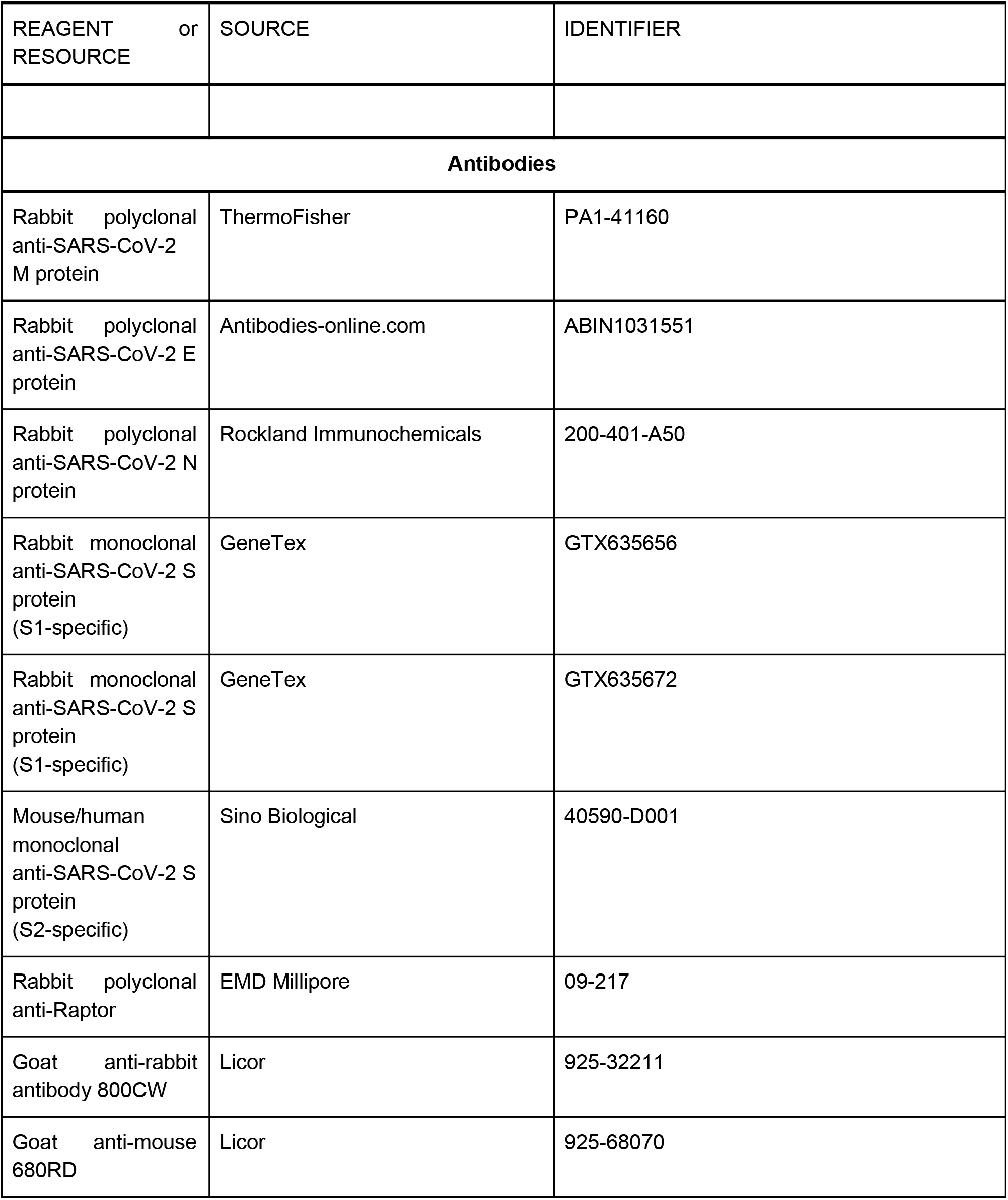
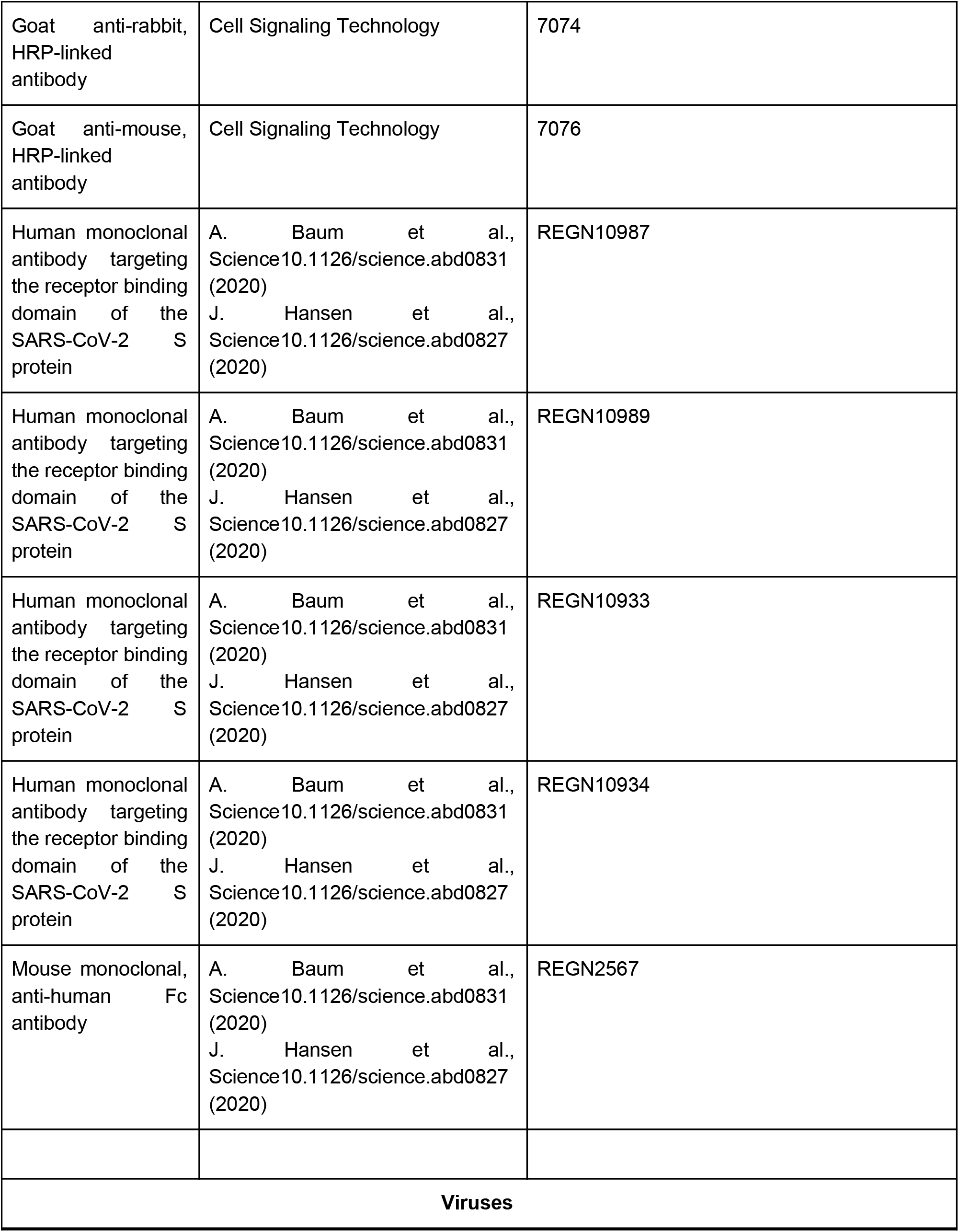
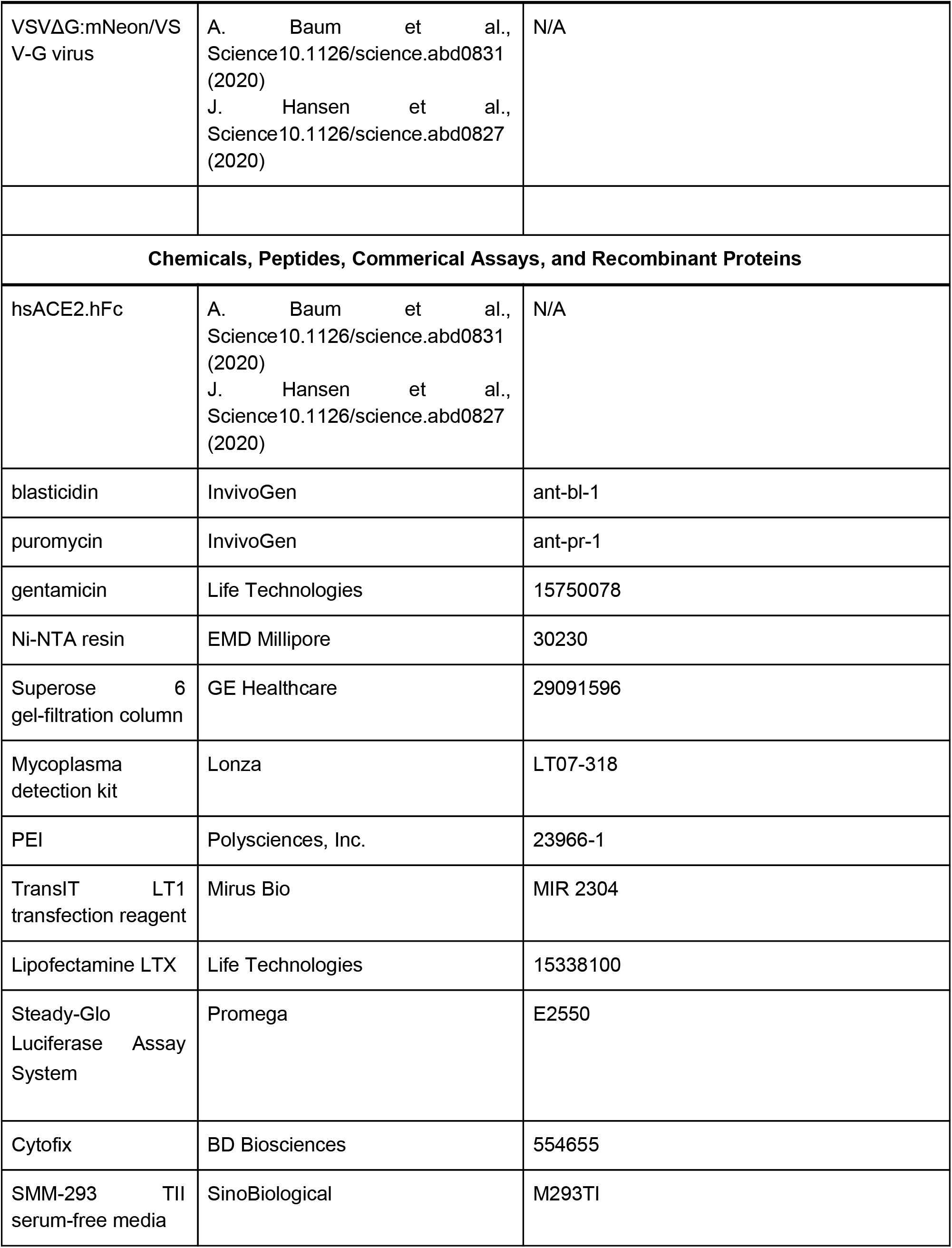
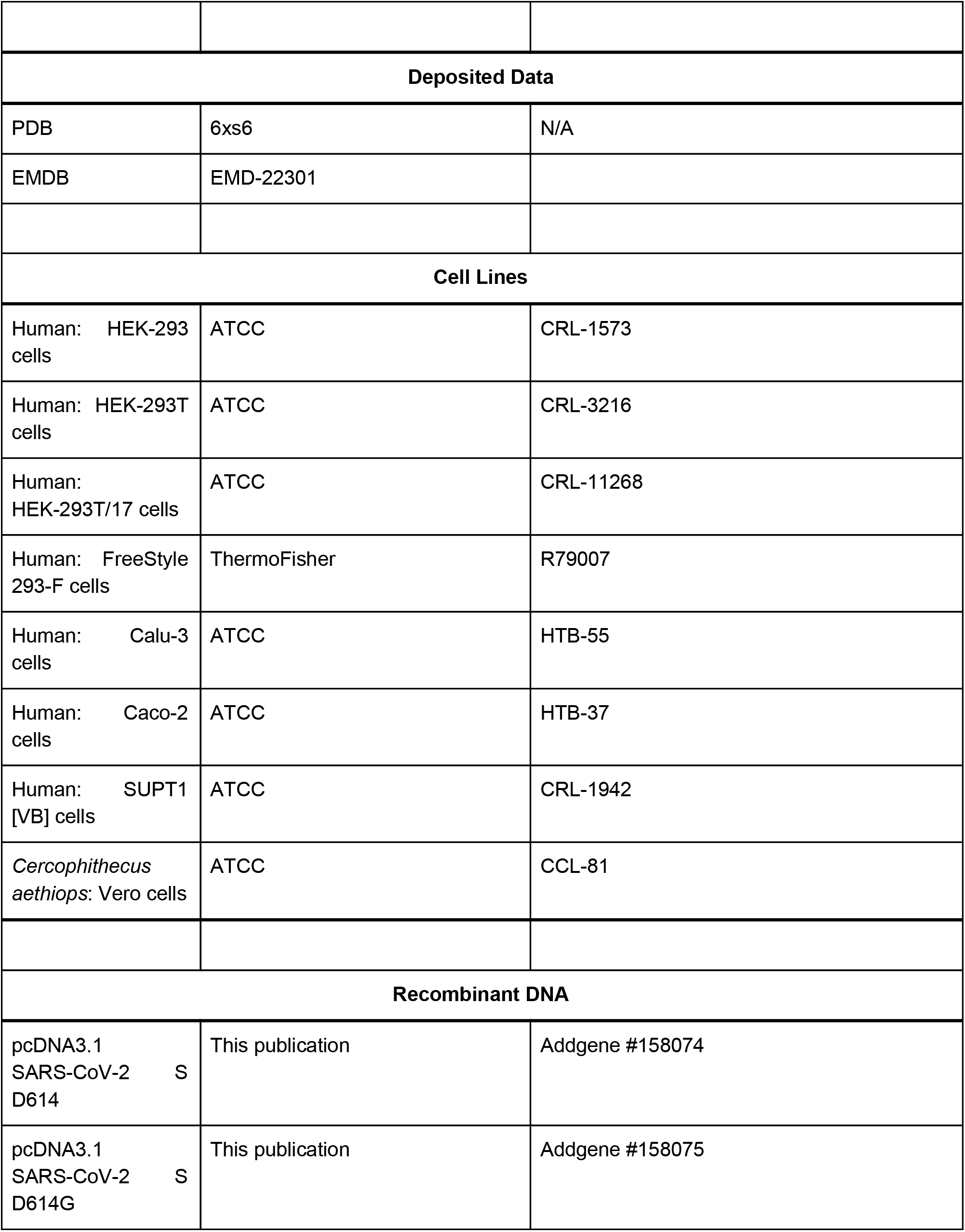
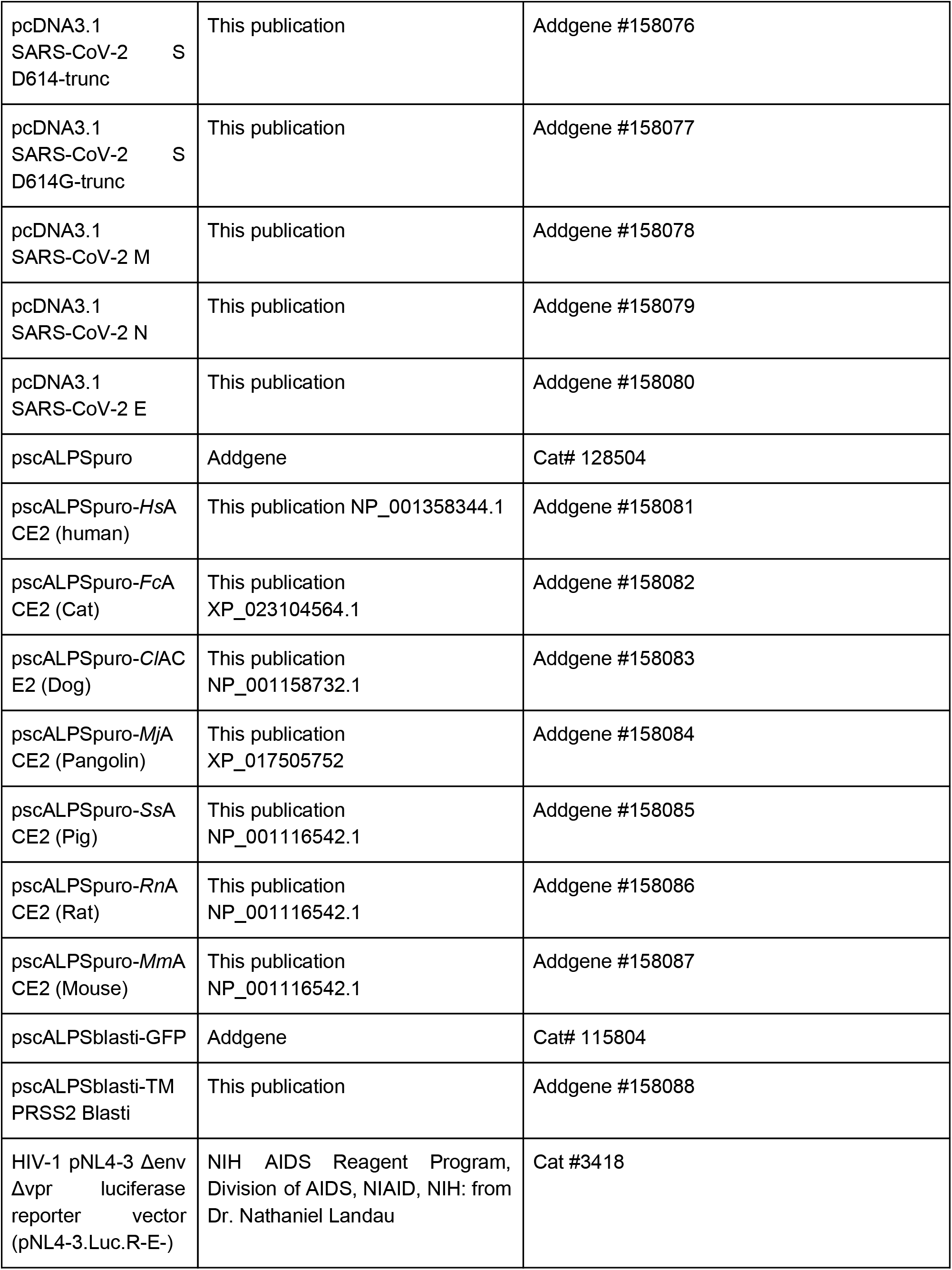
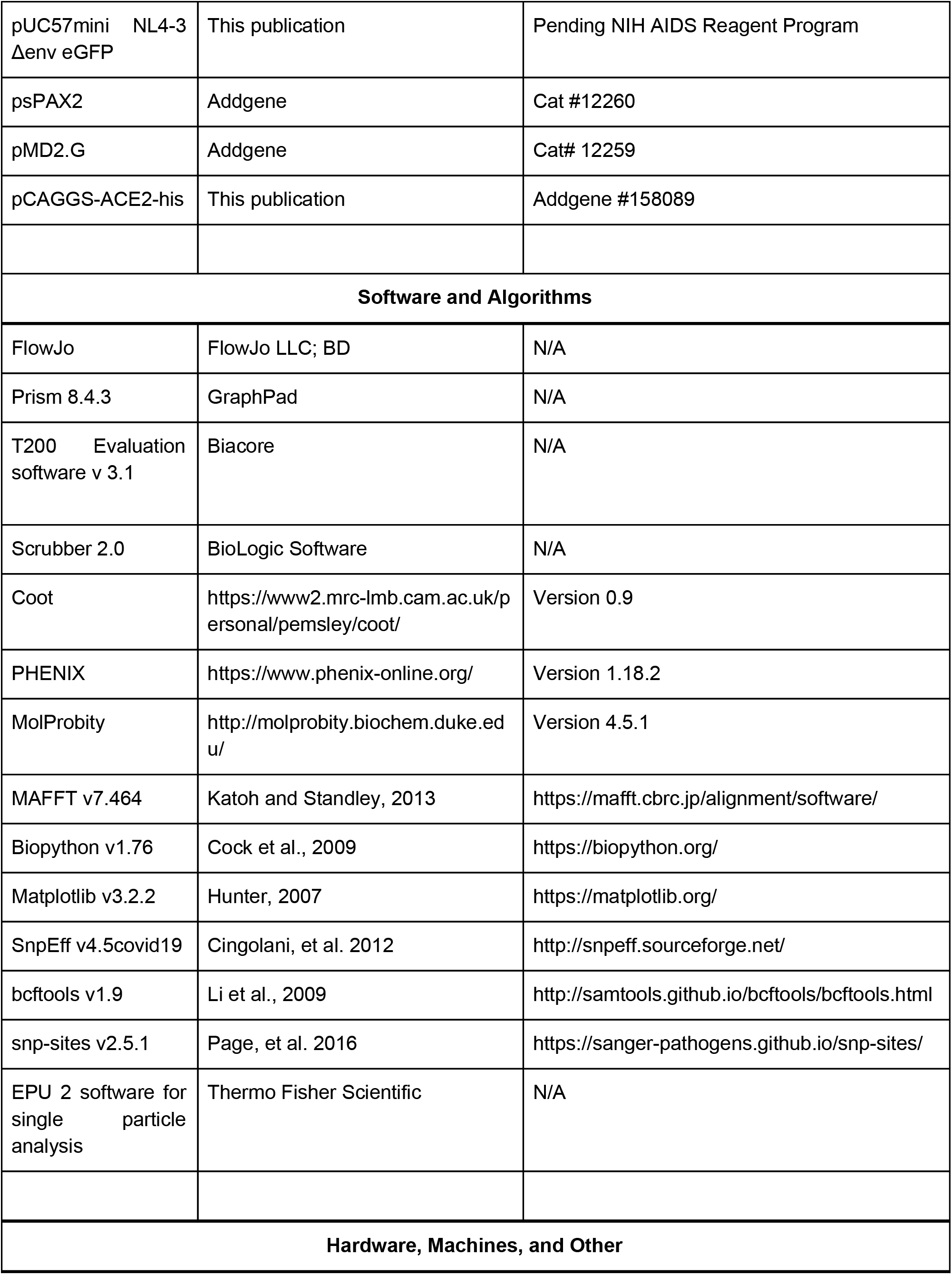
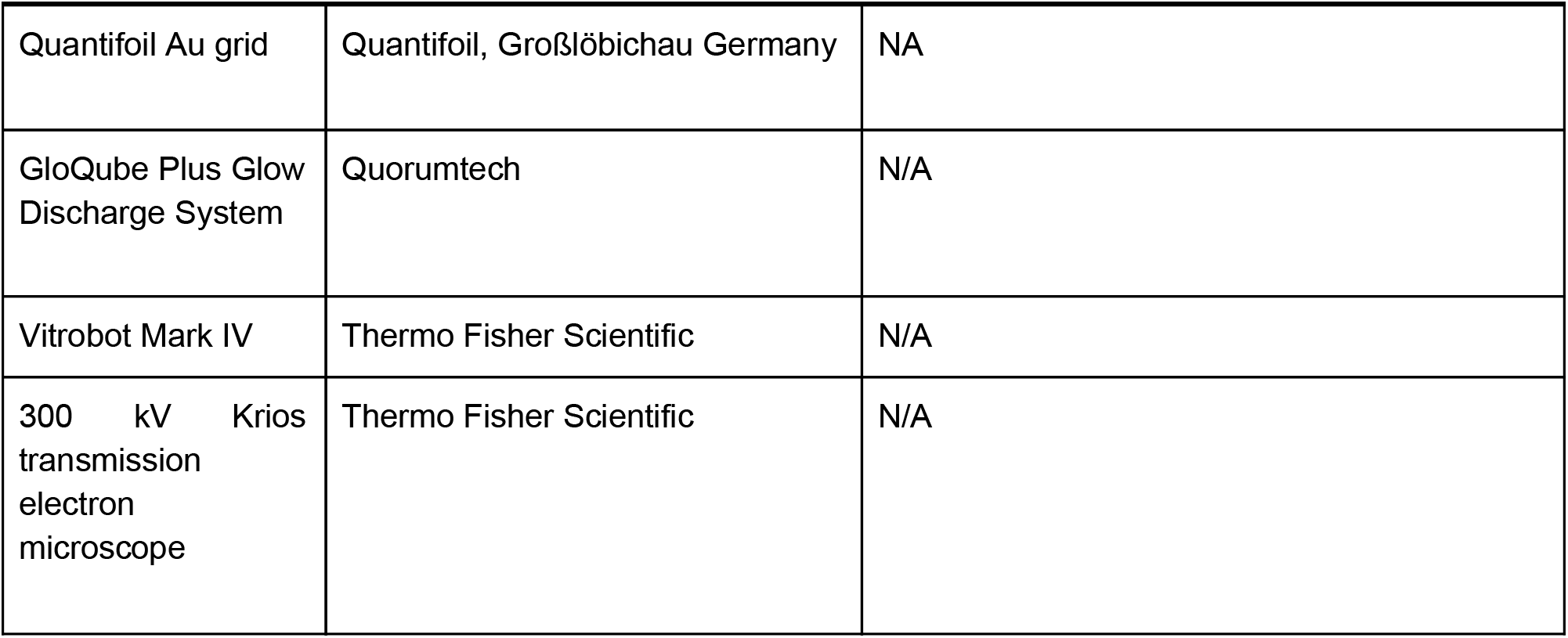
List of Reagents and Resources used in this manuscript

## Notes

### Summary of Updates

Data added includes: SPR binding kinetics and cryoEM structural data

